# The extended Price equation: migration and sex

**DOI:** 10.1101/2021.09.28.462138

**Authors:** Jake Brown, Jared M. Field

## Abstract

The Price equation provides a general partition of evolutionary change into two components. The first is usually thought to represent natural selection and the second, transmission bias. Here, we provide a new derivation of the generalised equation, which contains a largely ignored third term. Unlike the original Price equation, this extension can account for migration and mixed asexual and sexual reproduction. The notation used here expresses the generalised equation explicitly in terms of fitness, rendering this otherwise difficult third term more open to biological interpretation and use. This re-derivation also permits fundamental results, derived from the Price equation, to be more easily generalised. We take Hamilton’s rule as a case study, and provide an exact, total expression that allows for population structures like haplodiploidy. Our analysis, more generally, makes clear the previously hidden assumptions in similar fundamental results, highlighting the caution that must be taken when interpreting them.

## Introduction

A primary goal of evolutionary theory is to analyse how phenotypic change occurs over time. There are many forces that can cause such change: mutation, transgenerational plasticity and migration, to name a few. However, the key force of interest in most studies is natural selection. This is because it is the long-term driver of adaptation and gives direction to evolutionary change. It is therefore particularly useful to isolate its influence from all other forces.

One of the most useful tools to do this was developed by Price (1970, 1972a). The Price equation is an abstract partition of total evolutionary change into two terms. The first is intended to represent natural selection and the second accounts for any residual change that occurs during parent-offspring transmission of traits. These are commonly referred to as the natural selection and transmission bias terms respectively (Frank, 2012; Okasha, 2006; Gardner, 2020; Okasha & Otsuka, 2020). In most instances, parents and offspring are not clones of one another, so transmission bias will be non-zero. Despite this, the structure of the Price equation allows for the natural selection term to be analysed in isolation. Following this method, many historical results such as Hamilton’s rule and Fisher’s fundamental theorem have been rederived using the Price equation (Queller, 1992; Frank, 1997). Such results focus on natural selection alone but can be generalised to include the transmission bias term as well, giving a complete expression for total evolutionary change. These exact expressions can be useful for solving more complex evolutionary problems that involve forces beyond natural selection (Frank, 1997).

However, the standard form of the Price equation is not fully general to *all* biological situations. Price (1970) himself noted this in his original paper: ‘equation 1 fails if gene *A* ploidy is not the same in each *P*_1_ member.’ It is therefore well known that the Price equation does not account for migration or mixes of asexual and sexual reproduction. There have been several attempts to correct for this in the literature. Kerr and Godfrey-Smith (2009) did so by creating a connection-based partition and Grafen (2015) developed a class-structured Price equation that allowed for both sex and age-classes. Both of these approaches have their applications (for example, see Fox & Kerr, 2012). Yet these extended partitions cannot be easily reconciled with the fundamental theorems of evolution as derived from the Price equation (Queller, 2017).

In this paper, we aim to connect these fundamental theorems to the extended Price equation given by Kerr and Godfrey-Smith (2009). We do this by first rederiving the extended equation using a new method. This yields an expression that is identical in form to the original Price equation, but with an additional term. Second, we discuss what this additional term represents, how it can be used, and when it has an explicit causal interpretation (Okasha & Otsuka, 2020). We note that the extra term allows for the population structures of migration and sex to be integrated into the Price equation. Third, we use a standard quantitative genetic model to derive a genetical form of the extended equation. Using this equation, we follow a similar path to Frank (1997) to derive a total, exact expression for Hamilton’s rule (Hamilton, 1964). We highlight that this extended rule can be applied to situations of haplodiploidy, which was not previously possible with the standard Price equation. Fourth, we consider applications to modelling and derive a simple model for the evolution of altruism with migration. Here, the extended partition allows the modeller to identify and quantify the specific causes of change. Finally, we summarise our findings and consider future applications of our equations.

## Definitions

Consider an ancestral population *A* containing *n*_*A*_ ancestors and a descendant population *D* containing *n*_*D*_ descendants. Let the absolute fitness of an ancestor, *w*_*i*_, be given by the number of descendants that they have in *D*. In the language of parent-offspring relationships, we think of *w*_*i*_ as the number of children that the *i*th ancestor bears. Let 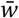 denote the average fitness of all ancestors in *A*. Additionally, we also define an index set *W*_*i*_, which contains the indices of all the descendants of the *i*th ancestor. Its cardinality is therefore *w*_*i*_.

In the descendant population *D*, we let *a*_*j*_ be the number of ancestors of a particular descendant *j*. Similar to fitness, we can think of *a*_*j*_ as the number of parents of the *j*th descendant. As such, we refer to *a*_*j*_ as the ancestry value of the *j*th descendant. Let *ā*_*D*_ be the average ancestry value of all descendants. Since we are now dealing with averages over two different sets, we use the subscript *D* to indicate that *ā*_*D*_ is an average over descendants not ancestors. Again, we also define the index set *A*_*j*_ which contains the indices of all the ancestors of the *j*th descendant and which has cardinality *a*_*j*_. The definitions above are summarised graphically in Figure 1.

**Figure 1:**
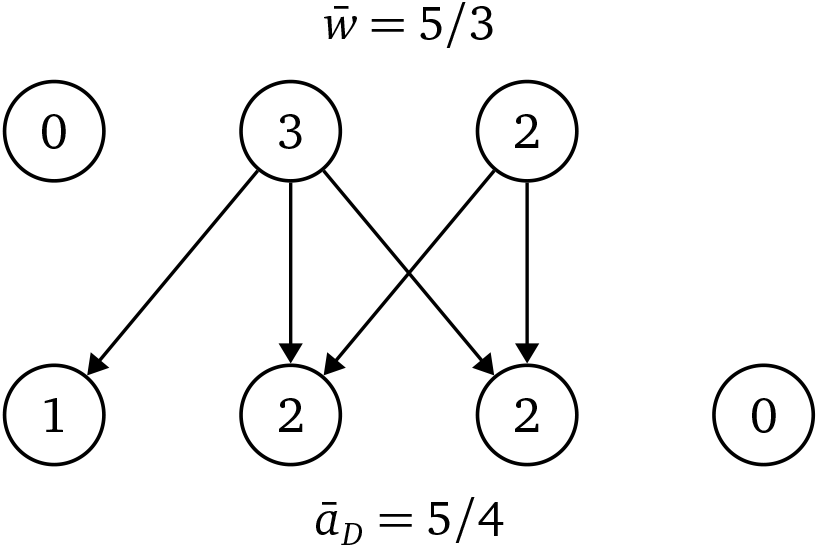
Definitions of fitness and ancestry value. Circles in the top row represent ancestors in *A* with each interior number indicating absolute fitness (*w*_*i*_). Circles in the bottom row represent descendants in *D* with each interior number indicating ancestry value (*a*_*j*_). Arrows are parent-offspring relationships.

We can now derive two identities which will be useful in the next section. First, the total number of connections (arrows) between *A* and *D* is:

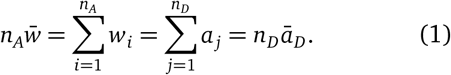

This is just two different ways of counting the total number of parent-offspring connections. Second, we can also replace *w*_*i*_ and *a*_*j*_ in Equation 1 above with summations over their corresponding index sets to get:

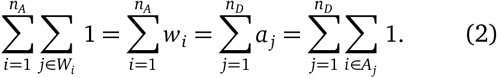

Here, the unit constant function under the summations is acting as a counter, similar to Kerr and Godfrey-Smith’s (2009) indicator variable. However since *n*_*D*_ and *n*_*A*_ are finite, we can replace this constant function with any arbitrary function *f*_*ij*_, and still interchange the order of summation according to Equation 2.

As is standard with the Price equation, eventually we aim to track some trait value *z* across the two populations. To this end, let an ancestor in *A* have trait value *z*_*i*_ and let their (potential) offspring have average trait value 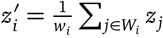. We use dashes to indicate a property of *D* as seen from the perspective of *A*. This feature of the standard Price equation will become significant later on. We can then let 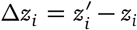 denote the average change in trait value between a parent and their offspring. Across all of *A* and *D*, we let 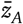 and 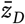 be the average trait values of the ancestor and descendant populations respectively. Finally, let 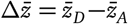 denote total average change between *A* and *D*.

## Derivation

Using these definitions, the next step in typical derivations of the Price equation is to make the following assumption (Frank, 2012; Gardner, 2020; Okasha, 2006):

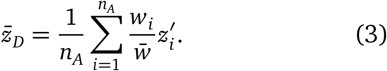

This assumption however, restricts several possible population structures, and is not fully general. For example, migration can affect the left hand side of Equation 3 by changing descendant average trait value, but it certainly cannot affect the right hand side, since migrants cannot be detected by 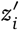 and *w*_*i*_ alone. To derive an extended partition, we will not make this assumption. Instead, we seek a remainder term by rearranging Equation 3 and simplifying using Equations 1 and 2:

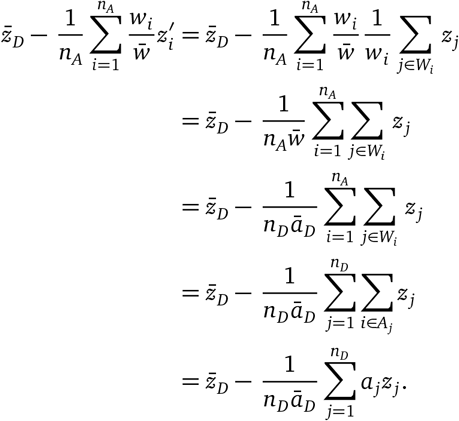

The second term is equal to E_*D*_(*az/ā*_*D*_) and since E_*D*_(*a/ā*_*D*_) = 1 we can write 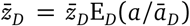. Using the statistical definition of (population) covariance then yields:

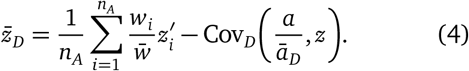

Equation 4 is fully general as opposed to Equation 3, so this highlights that the original Price equation assumes that Cov_*D*_(*a, z*) = 0. We discuss the significance of this below. To derive the extended partition, we can now follow any of the standard derivations of the Price equation listed above whilst using Equation 4 in place of Equation 3. In short, we do this by applying the identity 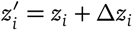 and then simplify using the statistical definitions of expectation and covariance to get:

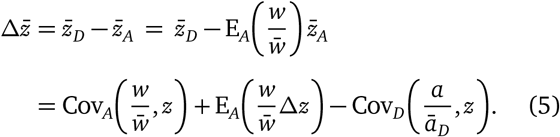

Equation 5 is written in terms of relative fitness 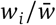 and relative ancestry value *a*_*j*_*/ā*_*D*_. For notational convenience, it is often easier to multiply both sides of Equation 5 by 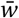 to remove fractions and deal with absolute fitness and ancestry value instead:

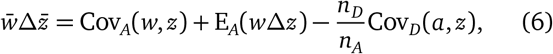

where 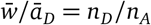 by Equation 1.

## Interpretation of the third term

How do we interpret Cov_*D*_(*a, z*)? At first the answer seems quite simple. This term measures how much the trait value of descendants (*z*) varies with the number of ancestors that each descendant possesses (*a*). Thus, we call this the *ancestry bias* term (Figure 2).

**Figure 2:**
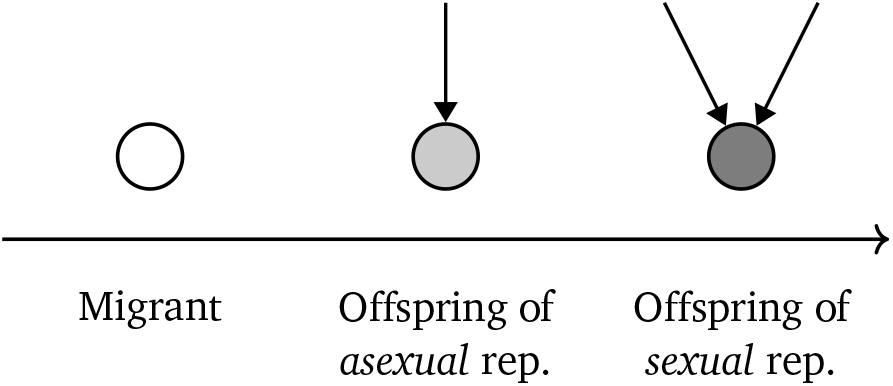
The ancestry bias term, Cov_*D*_(*a, z*)*/ā*_*D*_, detects whether there is a statistical association between ancestry value and trait value. Shading here represents different trait values for descendants in *D* and arrows are implied parental connections to *A*.

Using this straightforward covariance interpretation, we can quickly identify two biological cases where this term will be non-zero. First, in the case of migration where individuals from a population with a different average trait value, immigrate into *D*. Second, if ancestors reproduce both asexually and sexually, with offspring trait value dependent upon the mode of reproduction — reproductive bias. One interesting case of this are horizontal gene transfers (HGT) of antibiotic resistance genes between bacteria (von Wintersdorff et al., 2016). Bacteria that receive a HGT are more likely to receive a resistance gene and are therefore more likely to survive in the presence of the particular antibiotic in question. Whilst those bacteria that are only produced via binary fission, with no HGT, are less likely to carry the resistance gene and are therefore more likely to die. In terms of the internal book-keeping of the Price equation, these bacteria can be considered to have two parents. In this way, ancestry bias will be non-zero since ancestry value covaries with trait value.

These interpretations give an intuitive description of what the ancestry bias term describes, but what if we try to understand the covariance *causally*? There are three clear possibilities for causal relationships, shown in Figure 3. In the case of horizontal gene transfers above, ancestry value is influencing trait value (Figure 3b). Another example of this type of relationship is the sex determination of honey bees (or any haplodiploid species). Unfertilised eggs, the result of asexual reproduction, become male drones whilst fertilised eggs, the result of sexual reproduction, become female workers (Figure 4).

**Figure 3:**
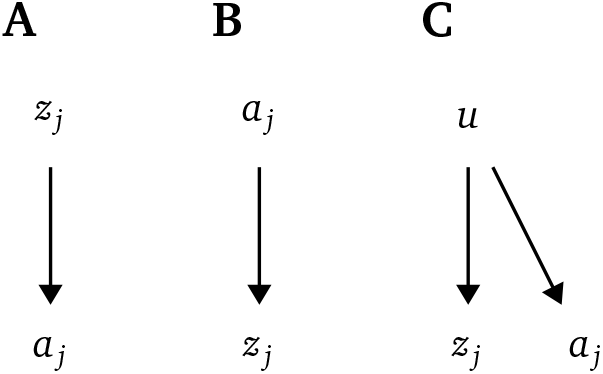
**(A)** Causal dependence of *a*_*j*_ on *z*_*j*_. **(B)** Causal dependence of *z*_*j*_ on *a*_*j*_. **(C)** An unidentified cause *u* that results in *z*_*j*_ and *a*_*j*_ being correlated.

**Figure 4:**
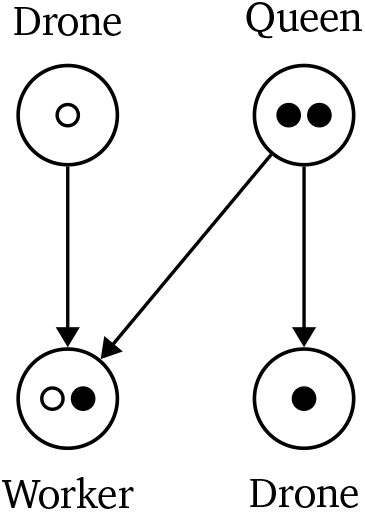
Multi-level selection in eusocial haplodiploid species such as honey bees. Larger circles are individuals, inner circles are alleles at a particular gene locus. Male drones are haploid, female workers and queens are diploid. Counting alleles, both the drone and the queen have a genetic fitness of 1 since there is one exact copy of their respective genotypes among descendants. However, the queen has an individual fitness of 2, whilst the drone has an individual fitness of 1. See Okasha (2006) for an in-depth review on multi-level selection.

On the other hand, trait value can also influence ancestry value (Figure 3a). This interpretation lends itself quite easily to cases of migration. For instance, we could consider a population with several distinct groups and migration between them. If faster individuals are more likely to migrate and slower individuals more likely to be philopatric, then trait value is clearly influencing ancestry value. On a genetic level, there are also genes that can predispose an individual to be migratory (Liedvogel et al., 2011). Further still, some other unidentified factor *u* might be influencing both descendant ancestry value and trait value causing Cov_*D*_(*a, z*) to be non-zero (Figure 3c).

This discussion highlights that we cannot simply assume what Cov_*D*_(*a, z*) represents *causally* by inspection (Okasha & Otsuka, 2020). In general, it is not even clear whether the ancestry bias term will always have an evolutionary interpretation. The key issue lies in the fact that migration and reproductive bias are both accounted for by the ancestry bias term, despite the fact that these are not typically seen as biologically related phenomena. Indeed, a priori, there are few reasons to suspect that migration and reproductive bias should be at all related. For this reason, if both of these phenomena are present in the model we are studying, this term is unlikely to have an interpretation beyond the statistical association shown in Figure 2.

It is possible however, to get clear causal partitions of the ancestry bias term when models are restricted slightly. For instance, when migration is present with only one mode of reproduction. In this case, migration will be the only force changing the ancestry bias term meaning that Cov_*D*_(*a, z*) *can* be interpreted as the effect of migration.

## Partial Fisherian change

Many applications of the Price equation seek to describe the part of total evolutionary change that is caused by natural selection, in a constant environment. First considered by Fisher (1930), this partial change became the focus of many subsequent evolutionary theorems, rules and equations (Queller, 2017). We call this partial Fisherian change and denote it by 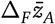 (Frank, 1997). In this section, we use the extended Price equation to derive an exact expression for total evolutionary change, from which we can more clearly interpret what exactly partial Fisherian change represents.

In classical quantitative genetics, the phenotype of a particular trait *A* is often written using the model:

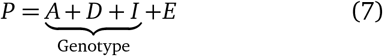

where *A* represents the additive genetic component of the phenotype, *D* is the effect of dominance, *I* the effect of epistasis between genes and *E* is the influence of the environment. Fisher (1930) was particularly interested in the heritable (additive) components of traits. In the model above, this is *A* but in the rest of the Price equation literature, it is usually written as *g*. We call *g* the additive genetic value or breeding value of a particular trait *z*. For a diploid organism, it is calculated as follows. Consider all possible alleles *ℓ* at every gene locus for a particular organism. Each organism will have *x*_*ℓ*_ = 0, 1 or 2 copies of allele *ℓ*. Using partial linear regression, we can then write:

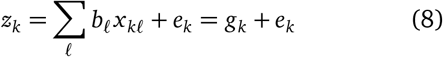

where *b*_*ℓ*_ is the average effect of allele *x*_*ℓ*_ on the phenotype. This linear model is fully general unlike Equation 7, which assumes independence of the right-hand side terms.

In practice, most of the average effects (*b*_*ℓ*_), will be close to zero meaning that we only need to index over a handful of alleles rather than all of them. In any case, by standard linear regression theory, Cov(*g*_*k*_, *e*_*k*_) = 0 so by definition, there is no interaction between *g*_*k*_ and *e*_*k*_. We also have E(*e*) = 0 which means that E(*z*) = E(*g*). We have intentionally chosen the index *k* above to be abstract as it applies to both populations *A* and *D*. Thus 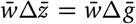. Substituting *z* = *g* into Equation 6 then yields:

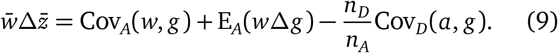

To derive the expression for partial Fisherian change now requires setting [E_*A*_(*w*Δ*g*)−Cov_*D*_(*a, z*)] → 0 to get:

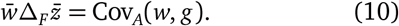

Equation 10 is known as Robertson’s secondary theorem of natural selection and it can be used to prove several other theorems: Fisher’s fundamental theorem, the general form of Hamilton’s rule (HRG) and the breeder’s equation (Frank, 1997, 2012). In a slightly different form, Fisher (1930) made the claim that Equation 10 represents the change in trait value that *would* occur if the environment were constant and if the only force acting upon the population were natural selection.

At face value, this claim seems reasonable if we assume that Cov_*A*_(*w, g*) is a faithful, causal representation of the effect of natural selection and if the other missing terms encapsulate all other effects. Setting the latter two terms of Equation 6 to zero is equivalent to isolating the force of natural selection in a constant environment and we are then left with Fisher’s interpretation.

There are a few problems with this line of reasoning. The key one is that Fisher’s interpretation of the environment was much broader than what most biologists would consider constitutes the environment today. Recall that *g* only represents additive genetic effects so dominance, epistasis and frequency dependence of alleles on others in the population are not encapsulated by Cov_*A*_(*w, g*). Under Fisher’s interpretation then, these effects are all part of the environment (Price, 1972b; Ewens, 1989; Okasha, 2008). This is quite different to the model shown in Equation 7.

Furthermore, many of the discussions about partial Fisherian change listed above are based on setting E_*A*_(*w*Δ*g*) → 0 or making equivalent assumptions. However, we now know that this condition is actually *not* sufficient to arrive at Equation 10, since there is the extra ancestry bias term to deal with. In addition to ignoring the effects of dominance, epistasis and frequency dependence, we also require setting Cov_*D*_(*a, g*) → 0. Note that we will ignore the case where Cov_*D*_(*a, g*) = E_*A*_(*w*Δ*g*) since this equality is unlikely to be stable.

For ancestry bias to be zero, we require the absence of all the possible effects described in the previous section. In most biological cases, it makes sense to ignore migration, as this is usually considered to be a distinct evolutionary force from natural selection. Excluding the other effects however, is more complex. This would set aside phenomena like haplodiploidy and horizontal gene transfers. These are not typically seen to be separate forces from natural selection; they are merely features of the reproductive structure of the species in question. Hence, the force of haplodiploidy is not typically investigated separately from natural selection. This is all to say that it is very natural to consider the change due to natural selection in a haplodiploid population. Yet such an analysis would require the ancestry bias term, or equivalent, and cannot be captured by Cov_*A*_(*w, g*) alone.

Equation 10 therefore describes change in populations where phenomena described by the ancestry bias term are absent. This is an important point, especially when considering social evolution, where hap lodiploidy is often of particular interest (Fromhage & Kokko, 2011; Gardner et al., 2012). This is primarily because many eusocial species have haplodiploid genetics. Analysis of such models with theorems that come from Equation 10, will ignore these features of population structure. For instance, HRG alone will not be able to distinguish between haplodiploid reproduction and standard diploid reproduction (Queller, 1985). This is somewhat concerning given that haplodiploidy was the initial inspiration for Hamilton (1964). However, we can easily correct for this issue by including both the transmission and the ancestry bias term in HRG. The result is an exact expression for total evolutionary change, whereby 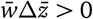 if and only if:

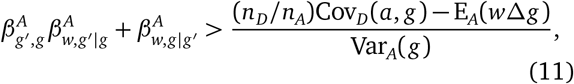

where each *β*^*A*^ is a regression coefficient over the population of ancestors and *g*′ is the average genetic value of social partners. See Queller (1992), Frank (1997) and Gardner et al. (2011) for full derivations and in-depth discussion. Substituting 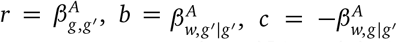 and *T* = [(*n*_*D*_*/n*_*A*_)Cov_*D*_(*a, g*) − *E*_*A*_(*w*Δ*g*)]/Var_*A*_(*g*) gives the familiar form of Hamilton’s rule with an additional term, allowing for effects like haplodiploidy: *r b* − *c* > *T*.

This is an exact, total result that applies to all evolutionary situations that can be modelled with the extended Price equation. As we have discussed in length above, these applications are much wider in scope than the standard Price equation. Although, in the particular case where *T* = 0, we recover the usual form of HRG. Here, it is important to note that this standard form of HRG emerges directly out of the natural selection term of the Price equation. Yet Equation 11 accounts for all forces of evolution, not just natural selection. On the one hand, this may be useful in certain applications, but on the other, it complicates the relatively simple *r b* − *c* > 0. This however, is a necessary complication if we want to study the effect of haplodiploidy, within the framework of the extended version of HRG. If the standard form of HRG is used alone, then the effects of haplodiploidy will be ignored. Ultimately, whether haplodiploidy should be considered together with, or apart from natural selection, is up to the theorist.

There are two further caveats to bear in mind when using Equation 11. First, each side of the inequality is not independent from the other. This is namely because the value of Var_*A*_(*g*) appears on both sides of the inequality; it is used in the calculation of *r* for instance. Second, the effect of haplodiploidy on *T* may not be straightforward to understand. For example, haplodiploidy can have an effect on both transmission and ancestry bias (Figure 4). Depending on how exactly this occurs, it may result in either a positive or a negative value of *T*. Hence, it can be misleading to draw immediate conclusions from the value of *T* alone. This is in contrast to the game theoretic version of Hamilton’s rule, denoted HRS for the special case (Birch, 2014). With HRS, the effect of haplodiploidy on altruism is clear because of the use of Wright’s coefficient of relatedness. This is the form of the rule that Hamilton (1964) originally used when describing social evolution in haplodiploid insect species. However, HRS does not generalise well and fails for non-additive payoffs, as well as for more complex cases (Queller, 1985; Nowak et al., 2010; van Veelen, 2009). Equation 11 however, generalises just as well as the extended Price equation, making it applicable to a very wide array of circumstances (Gardner et al., 2011).

Using a similar method to that shown in this section, we can also derive total expressions for any of the other fundamental theorems of evolution. Namely, we can start from Equation 9 and follow their respective derivations to arrive at the extended versions (Queller, 2017; Frank, 1997). We will not provide these expressions here.

## Partitioning evolutionary forces

Another application of the extended Price equation is partitioning the dynamics of a particular model into specific evolutionary forces. This of course can be done using other methods, but in this section, we construct a simple model for studying the evolution of altruism within the framework of the Price equation. The ancestry bias term in particular is useful in comparing the effects of migration against natural selection. This analysis could also be done with the equations given by Kerr and Godfrey-Smith (2009). However, our equations offer two advantages. First, our notation makes it slightly clearer what each term represents. Second, our equations are an extension of the standard Price equation. This means that any previous models constructed using a Price equation approach can easily be extended to include migration and reproductive bias by simply adding in the ancestry bias term.

For our model, we consider an asexually reproducing population with two types of individuals: selfish (*z* = 0) and altruistic (*z* = 1). In the *n*th generation, let there be *s*_*n*_ selfish individuals, *l*_*n*_ altruistic individuals and let *p*_*n*_ = *l*_*n*_/(*l*_*n*_ + *s*_*n*_) be the proportion of altruistic individuals. Both types have a base fitness of *w*_0_ = 1 and receive a fitness benefit of *p*_*n*_ *b* from all altruistic individuals — the greater the proportion of altruistic individuals the greater the benefit received. Let altruists suffer fitness cost *c* for conducting the altruistic act and assume that altruists can detect selfish individuals and actively repel them from the group. This is similar to how the immune system can detect and destroy selfish cancer cells (Loose & Van de Wiele, 2009). Accordingly, selfish individuals suffer fitness cost *p*_*n*_*d* — the greater the proportion of altruistic individuals, the stronger selfish individuals are repelled. With this setup, the fitness values of altruistic and selfish individuals in generation *n* are respectively:

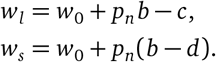

To prevent the overall population from becoming too large, we can adjust *w*_0_ to be smaller, so long as it remains positive. In doing this we must also be careful to avoid negative population sizes. This can be done by ensuring: *w*_*0*_ ≥ *d* − *b* and *w*_*0*_ ≥ *c*. For simplicity, assume that the altruistic trait is perfectly inherited such that transmission bias is zero. Using the standard Price equation we get (see Appendix for details):

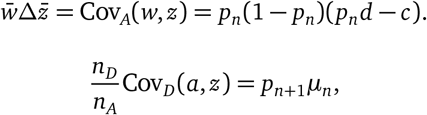

and therefore Equation 6 gives:

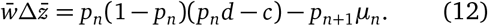

Equation 12 allows us both to model the dynamics of this toy model and to examine the causes of these dynamics (Figure 5). In our particular example, we see that the force of migration is relatively small whilst the population is large (*n* > 25). As the population shrinks due to the base fitness, *w*_0_ = 0.95, being less than 1, this force begins to increase. Then, as altruists become less frequent, selection switches to favour selfish behaviour rather than altruism.

**Figure 5:**
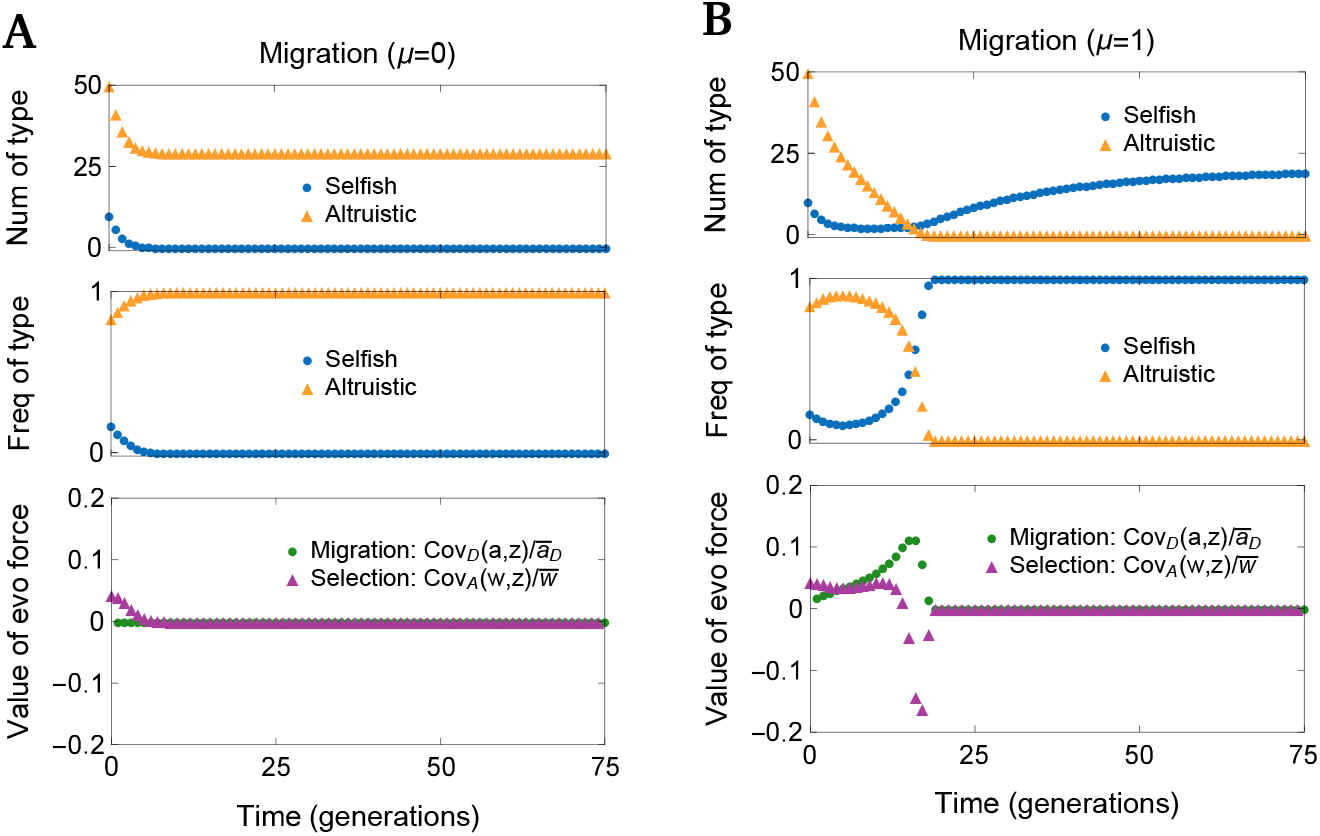
A simple model for the evolution of an altruistic trait. In both **(A)** and **(B)**, there are 50 altruistic individuals and 10 selfish individuals in generation 0. Base fitness is *w*_0_ = 0.95. Altruists provide fitness benefit *b* = 1.05 to all individuals in the population at fitness cost *c* = 1 for themselves. Selfish individuals are repelled from the population with strength *d* = 1.5. **(A)** The force of selection is always positive meaning that altruism is favoured by natural selection. This causes the altruistic trait to become rapidly fixed. **(B)** The force of migration is now added, with *µ* = 1 selfish individuals entering the population each generation. Altruism is initially favoured by natural selection, but as the population shrinks, the force of migration increases until selfishness becomes favourable. The altruistic trait is eventually lost, and the number of selfish individuals plateaus.

We could use this analysis to aid hypothetical experimental interventions to favour the evolution of altruism. For instance, to keep the force of migration small, it is ideal to keep the local population large. This is obvious in our simple case where all migrants are selfish. However, our model can also be extended to account for instances where only a proportion of migrants are selfish. Analysis like this is straightforward with the Price equation, since the partitioning of evolutionary forces is built into its structure, up to causal interpretation of the statistical terms. In this way, the extended partition expands the modelling capacities of the Price equation to match those achieved via other methods (Niehus et al., 2015).

## Discussion

To date, the standard Price equation has certainly been the best contender for the title of a fundamental theorem of evolution (Queller, 2017). Its abstract nature makes very few assumptions compared to almost every other equation in evolutionary biology. However, in this paper we have made clear the implicit assumptions of Price’s original equation. At a minimum, one of the four classical forces of evolution is missing: gene flow. Beyond this, the equation also assumes a constant mode of reproduction and equal ploidy amongst all individuals.

For this reason, applications of the Price equation have nearly always made simplifying assumptions such as: (1) excluding migration and (2) assuming only one mode of reproduction (exclusively asexual or sexual). These assumptions ensure that Cov_*D*_(*a, z*) → 0, and allow the standard Price equation to be used. It is also very common to make such assumptions when constructing models in population genetics. In light of this, it is not so surprising that the ancestry bias term has been largely neglected.

Despite this, it is still unclear what exactly goes wrong with the original two terms to fail to account for the expanded scenarios discussed above. The key to investigating this is noting that both the natural selection and transmission bias terms index over the ancestral population *A* alone (Frank, 1998; Okasha, 2006). Yet some aspects of population structure can only be detected by also indexing across *D* (Figure 6). Migration is one such example. If we only index over *A*, it will be impossible to find a migrant in *D*, who by definition has no connections to ancestors in *A* (Figure 6a, b).

**Figure 6:**
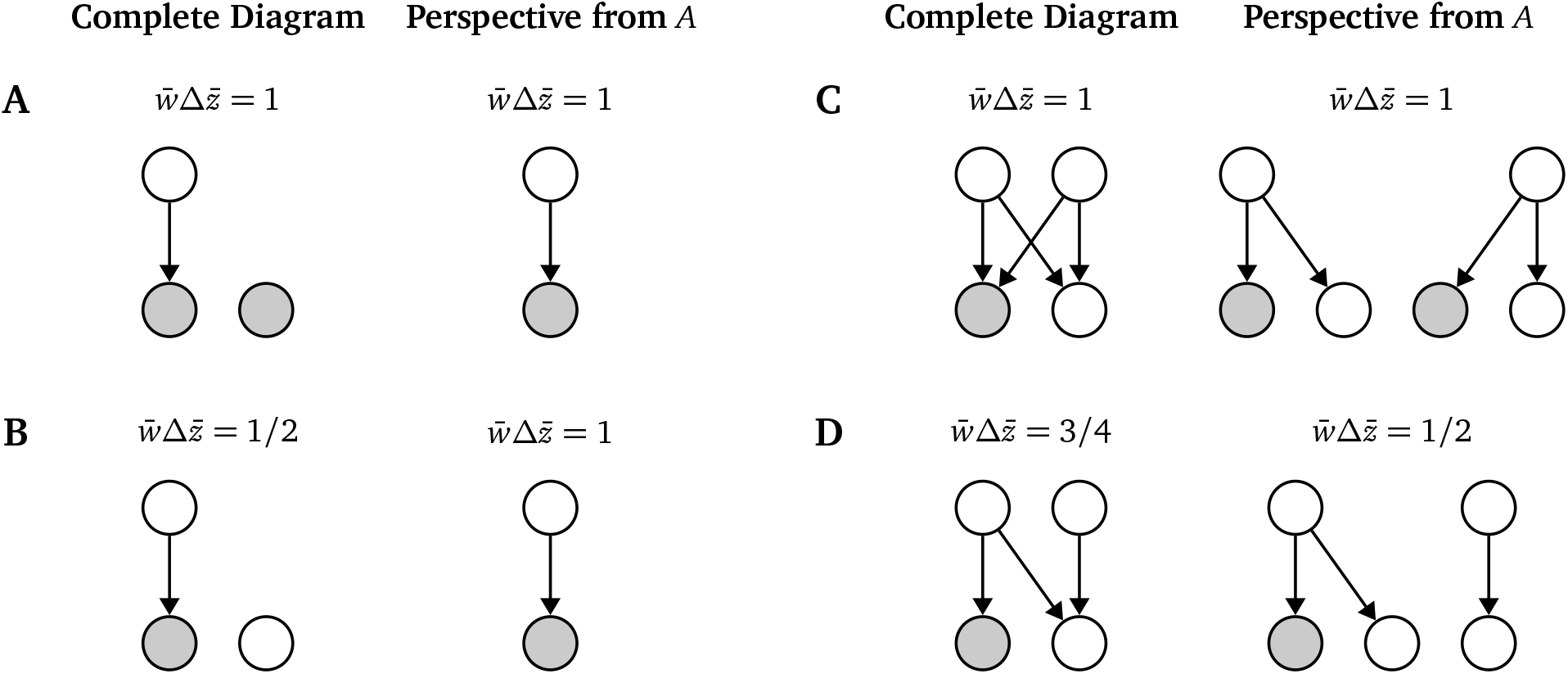
Evolutionary scenarios from complete and restricted perspectives. Arrows indicate parent-offspring relationships. Shaded individuals have trait value 1 and unshaded individuals have trait value 0. The perspective from *A* is the evolution perceived by natural selection, Cov_*A*_(*w, z*), and transmission bias, E_*A*_(*w*Δ*z*), alone. 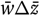 is the total evolutionary change as seen from each perspective. **(A)** Homogeneous migration where migrants appear the same as direct descendants. **(B)** Non-homogeneous migration where migrants introduce new variation. This is invisible from the perspective of *A*. **(C)** Completely sexual reproduction. From the perspective of *A*, there are twice the number of descendants, but the total evolutionary change is the same. **(D)** Mixed asexual and sexual reproduction. The perspective from *A* is inaccurate and results in an incorrect calculation of total evolutionary change.

In the case of mixed modes of reproduction, a miscounting problem can arise. To see how this occurs, first consider the case of constant sexual reproduction with two sexes (Figure 6c). Here, two parents reproduce sexually to have two children. Each parent has fitness *w*_*i*_ = 2 but from the perspective of *A*, this means quite bizarrely that there are apparently four children in total. However, this is not a problem since each descendant is counted exactly twice when indexing over *A*, resulting in the correct calculation of total evolutionary change 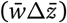. By contrast, if there is a mix of reproduction methods, then some descendants may be counted twice and others only once (Figure 6d). This can lead to a miscalculation of total evolutionary change.

Therefore, the view of evolution from the perspective of the ancestral population alone, is restricted, whilst the extended partition allows for a complete view of evolution. In situations where the perspective from *A* is incomplete, the ancestry bias term perfectly adjusts for this, giving the correct calculation of total evolutionary change 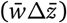. Thus, we can think of ancestry bias as a correction term.

## Conclusion

The extended Price equation provides an integrated approach to allow for a range of evolutionary phenomena, including migration and mixed asexual and sexual reproduction. These effects are all accounted for by a single extra term, Cov_*D*_(*a, z*), which represents ancestry bias. This term detects the covariance between a descendant’s trait value and the number of parents they have in the ancestral population. When a specific causal model is imposed, ancestry bias can also quantify particular evolutionary forces like migration. More generally, ancestry bias allows for mixed ploidies within the framework of the Price equation. With this extension, we can therefore connect phenomena like haplodiploidy to many standard fundamental theorems of evolution that are derived from the Price equation. For instance, a generalised form of Hamilton’s rule can be derived that accounts for haplodiploidy and migration.

There are also applications to modelling. The extended partition allows for the integration of migration and mixed modes of reproduction into Price-equation based models. This increases the domain of applicability of the Price equation, and may make it a suitable alternative to ODE-based models.

There are several further applications of the generalised Price equation that could be considered. First, asexual reproduction could be reinterpreted as a continuation of generations alongside standard sexual reproduction. Under this interpretation, simple models for overlapping generations could be constructed, offering an alternative to class-based models (Grafen, 2015). We envisage that this approach could be used to study life-history traits and social evolution between generations, such as in the Grandmother hypothesis (Cant & Johnstone, 2008). Second, applications to other fundamental theorems could be considered. In this paper we have focused on Hamilton’s rule, but the extended partition could easily be used to derive total expressions for Fisher’s fundamental theorem of natural selection or the breeder’s equation. Third, a multi-level selection (MLS) partition could also be imposed upon the extended Price equation. In fact, the standard derivation used to derive the MLS partition could also be applied to the ancestry bias term (Okasha, 2006; Gardner, 2015). This would allow for the study of intra-group versus inter-group ancestry bias. Finally, deeper causal analysis of the ancestry bias term using the counterfactual method employed most recently by Okasha and Otsuka (2020) would be insightful.

## Author Contributions

JB conducted the research and wrote the manuscript in consultation with JMF. JMF supervised the research.

## Acknowledgements

JB was funded by the Mathematics and Statistics Vacation Scholarships Program at the University of Melbourne. JMF is funded by a McKenzie Fellowship at the University of Melbourne.

## Appendix

### Derivation of altruism model

We use a set of tables to calculate Price equation terms, similar to Gardner et al. (2011).

The average trait value in *A* will simply be the proportion of altruists: 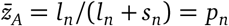. Multiplying across the rows in Table 1 and adding the appropriate results yields:

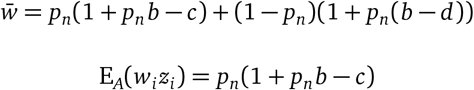

which means that the natural selection term is:

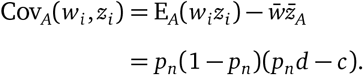

**Table 1:** Population data over *A*

| Fitness ($w_i$) | Trait value ($z_i$) | Frequency |
| --- | --- | --- |
| $1 + p_n b - c$ | 1 | $\frac{l_n}{l_n + s_n} = p_n$ |
| $1 + p_n(b - d)$ | 0 | $\frac{s_n}{l_n + s_n} = 1 - p_n$ |

Similarly, the average trait value in *D* will be the proportion of altruists: 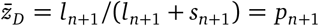. Again, we multiply across the rows in Table 2 and add the corresponding results together to get:

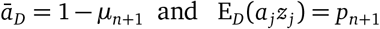

**Table 2:** Population data over *D*

| Ancestry value ($a_j$) | Trait value ($z_j$) | Frequency |
| --- | --- | --- |
| 1 | 1 | $\frac{l_{n+1}}{l_{n+1} + s_{n+1}} = p_{n+1}$ |
| 1 | 0 | $\frac{s_{n+1} - m}{l_{n+1} + s_{n+1}} = 1 - p_{n+1} - \mu_{n+1}$ |
| 0 | 0 | $\frac{m}{l_{n+1} + s_{n+1}} = \mu_{n+1}$ |

So the ancestry bias term is:

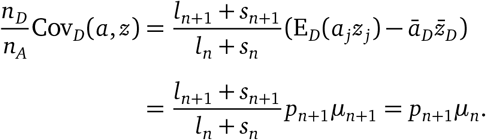

